# Pacbio sequencing reveals identical organelle genomes between American cranberry (*Vaccinium macrocarpon* Ait.) and a wild relative

**DOI:** 10.1101/567925

**Authors:** Luis Diaz-Garcia, Lorraine Rodriguez-Bonilla, Tyler Smith, Juan Zalapa

## Abstract

Breeding efforts in the American cranberry (*Vaccinium macrocarpon A*it.), a North American perennial fruit crop of great importance, have been hampered by the limited genetic and phenotypic variability observed among cultivars and experimental materials. Most of the cultivars commercially used by cranberry growers today were derived from a few wild accessions bred in the 1950s. In different crops, wild germplasm has been used as an important genetic resource to incorporate novel traits and increase the phenotypic diversity of breeding materials. *Vaccinium microcarpum* (Turcz. ex Rupr.) Schmalh. and *V. oxycoccos* L., two closely related species, may be cross-compatible with the American cranberry and could be useful to improve fruit quality such as phytochemical content, and given their northern distribution, could also help develop cold hardy cultivars. Although these species have previously been analyzed in diversity studies, genomic characterization and comparative studies are still lacking. In this study, we sequenced and assembled the organelle genomes of the cultivated American cranberry and its wild relative, *V. microcarpum*. PacBio sequencing technology allowed us to assemble both mitochondrial and plastid genomes at very high coverage and in a single circular scaffold. A comparative analysis revealed that the mitochondrial genome sequences were identical between both species and that the plastids presented only two synonymous single nucleotide polymorphisms (SNPs). Moreover, Illumina resequencing of additional accessions of *V. microcarpum* and *V. oxycoccos* revealed high genetic variation in both species. Based on these results, we provided a hypothesis involving the extension and dynamics of the last glaciation period in North America, and how this could have shaped the distribution and dispersal of *V. microcarpum*. Finally, we provided important data regarding the polyploid origin of *V. oxycoccos*.

## 1. Introduction

Worldwide the majority of food crops derive from a few wild plant species. In the US, farmers grow a great variety of plants such as cereals, sugar crops, vegetable and fruits (FAO, 2010). However, few crop species are native to North America and even fewer can still be found in their wild forms and habitats. Wild germplasm, especially from crop wild relatives, is of tremendous importance for breeding and agricultural production since it represents a pool of useful characteristics such as tolerance to biotic and abiotic stress, resistance genes, and traits influencing yield [1–4].

*Vaccinium macrocarpon* Ait. (American cranberry) is endemic to North America, and one of the few fruit crops that can still be found in the wild. The niche of this species is similar to its wild relative *Vaccinium oxycoccos* L. (small cranberry) [5–7]. However, *V. oxycoccos* is restricted to peatlands, where it grows embedded in moss lawns (usually sphagnum). *Vaccinium macrocarpon* grows in a wider variety of wetlands, including peatlands, swamps and wet shores, and it will grow in mineral soils as well as moss lawns. *Vaccinium oxycoccos* produces over-wintering berries that, although smaller than the cultivated cranberry, share a similar flavor and possess a superior antioxidant content [8,9]. The high phytochemical content of the wild relative species and the relatively low genetic diversity in cultivated cranberry, make the wild species extremely useful for breeding purposes [10,11]. In addition to its unique antioxidant profile, its northern and circumboreal distribution [12,13], make the small cranberry an interesting source of traits such as cold hardiness [14,15].

While the American cranberry is a diploid species, the small cranberry *V. oxycoccos* can be found as diploid, tetraploid and, rarely, hexaploid forms. Although the most recent Flora of North America treatment [6] treats all three cytotypes as a single variable species (i.e., *V. oxycoccos*), morphological, molecular, phenological and ecological data support recognition of the diploid form as a distinct species, *Vaccinium microcarpum* (Turcz. ex Rupr.) Schmalh. [11,16,17].

Camp (1944) considered *V. macrocarpon* the ancestral form, from which *V. microcarpum* was derived. According to this theory, the tetraploid *V. oxycoccos* formed following secondary contact of the diploid species, through a combination of polyploidy and hybridization. He noted that this could have followed two paths: allopolyploid hybridization between *V. macrocarpon* and *V. microcarpum,* or the hybridization of autotetraploid plants derived from each diploid species. Regardless of the exact mechanism, he considered the morphology of *V. oxycoccos,* which is in some ways intermediate between the two diploids, strong evidence supporting hybrid ancestry. This hypothesis is supported by allozyme data [19], which shows that *V. oxycoccos* possesses alleles from both of the diploid species. However, the same data revealed no fixed heterozygous loci, which is consistent with tetrasomic inheritance as expected in autotetraploids [19,20]. Our own AFLP data [11] also reveal the presence of alleles from both diploids in *V. oxycoccos,* although simulation analysis suggests that it has diverged from the diploids since its formation.

During the last decade, several studies regarding cranberry genetics and genomics have been published. Georgi and collaborators (2013) published the first SSR-based genetic map and reported several quantitative trait loci (QTL) for yield related traits and important phytochemicals. Subsequently, high-density genetic maps as well as more multi-trait QTL studies became available [22–27]. In addition, nuclear [28], plastid [29], and mitochondrial [30] genomes have been published recently. Regarding diversity between *V. macrocarpon* and wild relatives, most of the studies have been based on simple sequence repeat (SSR) markers. Zhu et al. (2012) developed and tested a set of nuclear SSR markers and further validated them using diverse samples including wild and cultivated cranberries. Similarly, Zalapa et al. (2015) studied genetic variability among wild *V. macrocarpon* populations, and found a rich clonal diversity within the species. In addition, the same authors were able to clearly differentiate, based on SSR markers, *V. microcarpum* and *V. oxycoccos* from *V. macrocarpon*, although some genotypes presented certain overlap between species. Additionally, Schlautman and collaborators, in two different studies, [33,34] developed 697 SSRs and 16-multiplexing panels containing 61 SSR markers, respectively and showed that they could be used to reliable differentiate cranberry genotypes. Conversely, Schlautman et al. (2017, [35]) identified 54 organelle-based SSR markers, and after testing in several diverse cranberry genotypes discovered no variation among the evaluated materials.

Given the recent advances on next-generation sequencing technologies, we generated Pacbio complete plastid and mitochondrial genomes of an Alaskan accession of *V. microcarpum* as well as updated the American cranberry organelle genomes. A comprehensive comparison between the organelles of these two species revealed near identical sequences, which indicates common ancestry with implications about the origin of cultivated *V. macrocarpon* in North America. Moreover, Illumina resequencing in *V. microcarpum* samples collected in eastern Canada suggested a process of genetic diversification. Finally, we provided more evidence regarding the polyploid origin of *V. oxycoccos*.

## 2. Materials and Methods

### 2.1. Plant material

For *V. macrocarpon*, we used the commercial cultivar ‘Stevens’, which was derived from a cross between two wild selections, ‘McFarlin’ (from Massachusetts, USA) and ‘Potter’s Favorite’ (from Wisconsin, USA). For *V. microcarpum*, we used an accession collected by N. Vorsa (Rutgers University) in southern Alaska and described in Mahy et al. (2000). Both plants were maintained clonally under greenhouse conditions at the University of Wisconsin-Madison (Wisconsin, USA). These accessions have been extensively used in different marker studies, providing a unique genetic fingerprint based on nuclear and organellar information as well as known genetic relationships based on relatively diverse samples of *V. macrocarpon, V. microcarpum,* and *V. oxycoccos* [19,31–33]. In Figure 1, we show distinctive plant characteristics and attributes of each our *V. microcarpum* and *V. macrocarpon* accession.

**Figure 1.**
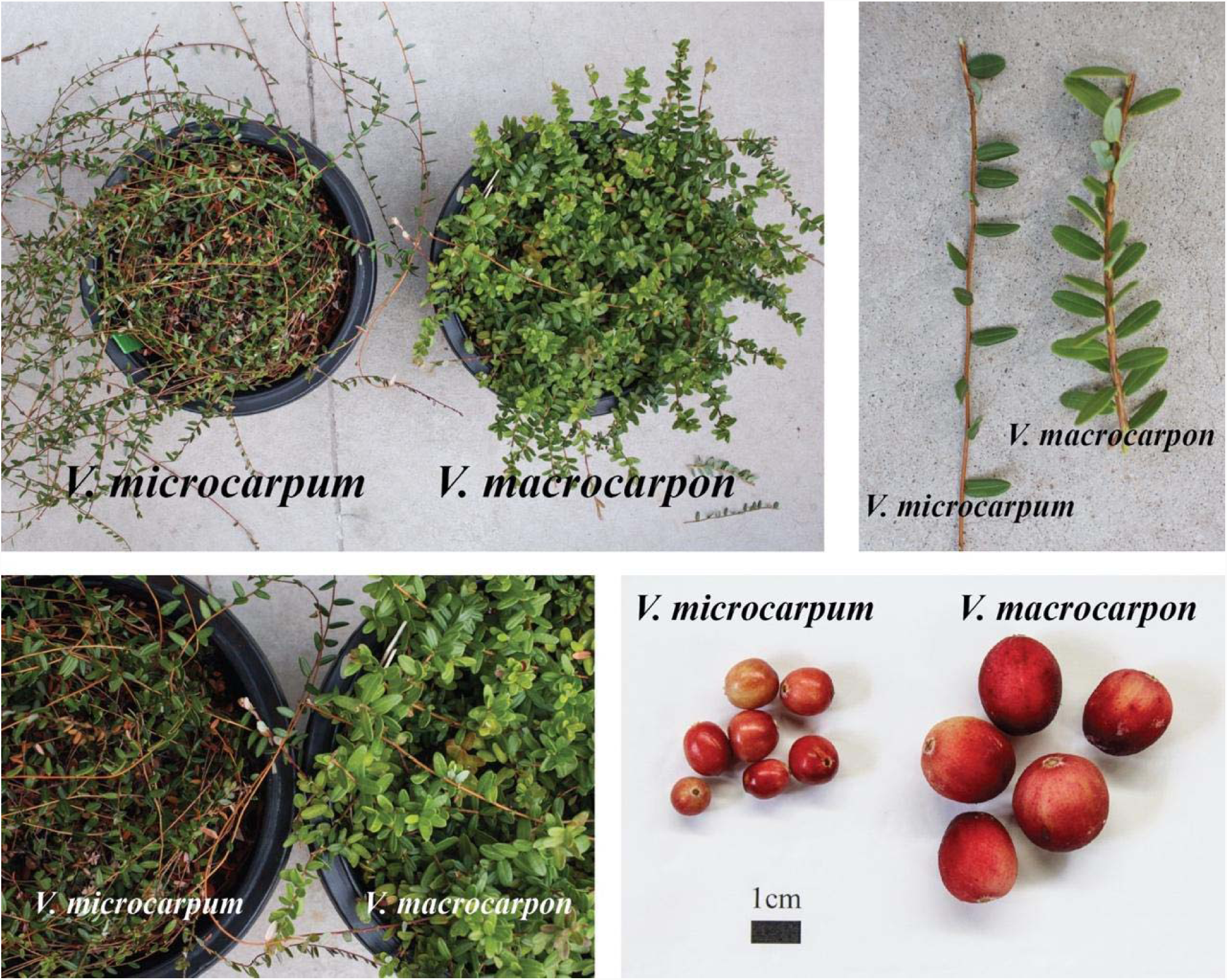
Distinctive characteristics of *V. microcarpum* and *V. macrocarpon* ‘Stevens’ plants.

### 2.2. Pacbio sequencing, genome assembly, and annotation

Young leaves of *V. macrocarpon* and *V. microcarpum* were processed at Amplicon Express (Pullman, Washington, USA) to obtain high molecular weight DNA. Two SMRT cells of PacBio Sequel were sequenced for each species at The DNA Technologies and Expression Analysis Cores of the University of California-Davis (San Diego, California, USA). For each species, a complete de novo assembly (including all the raw reads) was performed on Canu v1.7 [36] using the automatic pipeline (that includes read correction, trimming and assemble) with default parameters. From these assemblies, mapping (using blastn) the plastid assembly published by Fajardo et al. 2013 allowed us to extract contigs of lengths 209,726bp (for *V. macrocarpon*) and 223,859bp (for *V. microcarpum*), which we believed contain the complete plastid genomes. On the other hand, mapping the reference mitochondrial genome published by Fajardo et al. (2014) yielded contigs with lengths smaller than the expected genome size (459,678bp), therefore, an alternative strategy was carried out as follows. Raw PacBio reads were mapped with blasR (with -minMatch=15, and -fastSDP and -advanceHalf activated) using as reference the mitochondrial genome published by Fajardo et al. (2014); filtered reads were then de novo assembled using the automatic pipeline of Canu as in the assembling process used for the plastid. For *V. macrocarpon* and *V. microcarpum*, the largest contig on each assembly was 468,773bp and 468,164bp, respectively. Since Canu was ran using the default parameters, all assemblies were resolved at ∼40X coverage. As described in the user’s manual, Canu processes the longer reads up to get 40X, which are further corrected using the remaining shorter reads. Thus, a conventional estimation of coverage (which is computed using all the reads) would be considerably larger than this. A round of polishing was performed in the plastid and mitochondrial assemblies of both species using arrow (with default parameters). Subsequently, the circularity of each assembly was checked with the tool circlator v1.5.5 using -- b2r_length_cutoff=100000 and --merge_reassemble_end=9000. Following the recommendations of circlator’s authors, a final round of polishing with arrow was performed in all four assemblies, (i.e., two organelles each for *V. macrocarpon* and *V. microcarpum*).

Gene annotation was performed with GeSeq [37] using all the Ericales and Asterids as reference; the percentage of identity for the BLAT protein search, and for the BLAT rRNA, tRNA, and DNA search, were set to 70 and 80, respectively. All annotation were manually reviewed and corrected, if necessary, in Geneious R11 [38]. GC content was analyzed in R [39] using the package Biostrings. SSR search was performed with Phobos version 3.3.12 [40].

### 2.3. Illumina sequencing and mapping

We extended our study by sequencing additional accessions of *V. macrocarpon* and *V. microcarpum*, as well the tetraploid species *V. oxycoccos*. For *V. macrocarpon*, we included samples WABL11, WC16-13 (both wild accessions collected in Wisconsin, USA), and WC16-16, (wild accession collected in New Jersey, USA). For *V. microcarpum*, we include samples QCCW5, QCJB20 and QCWA9, all wild collected in Quebec, Canada (previously described in Smith et al., 2015). For *V. oxycoccos*, we include three wild samples, MWB3, PMS5 (both from Wisconsin, USA) and PB17 (from Minnesota, USA). Additionally, the *V. macrocarpon* and *V. microcarpum* samples used for Pacbio sequencing were also sequenced with Illumina. In Table 1, a summary of the samples used for both Pacbio and Illumina sequencing is provided.

**Table 1.**
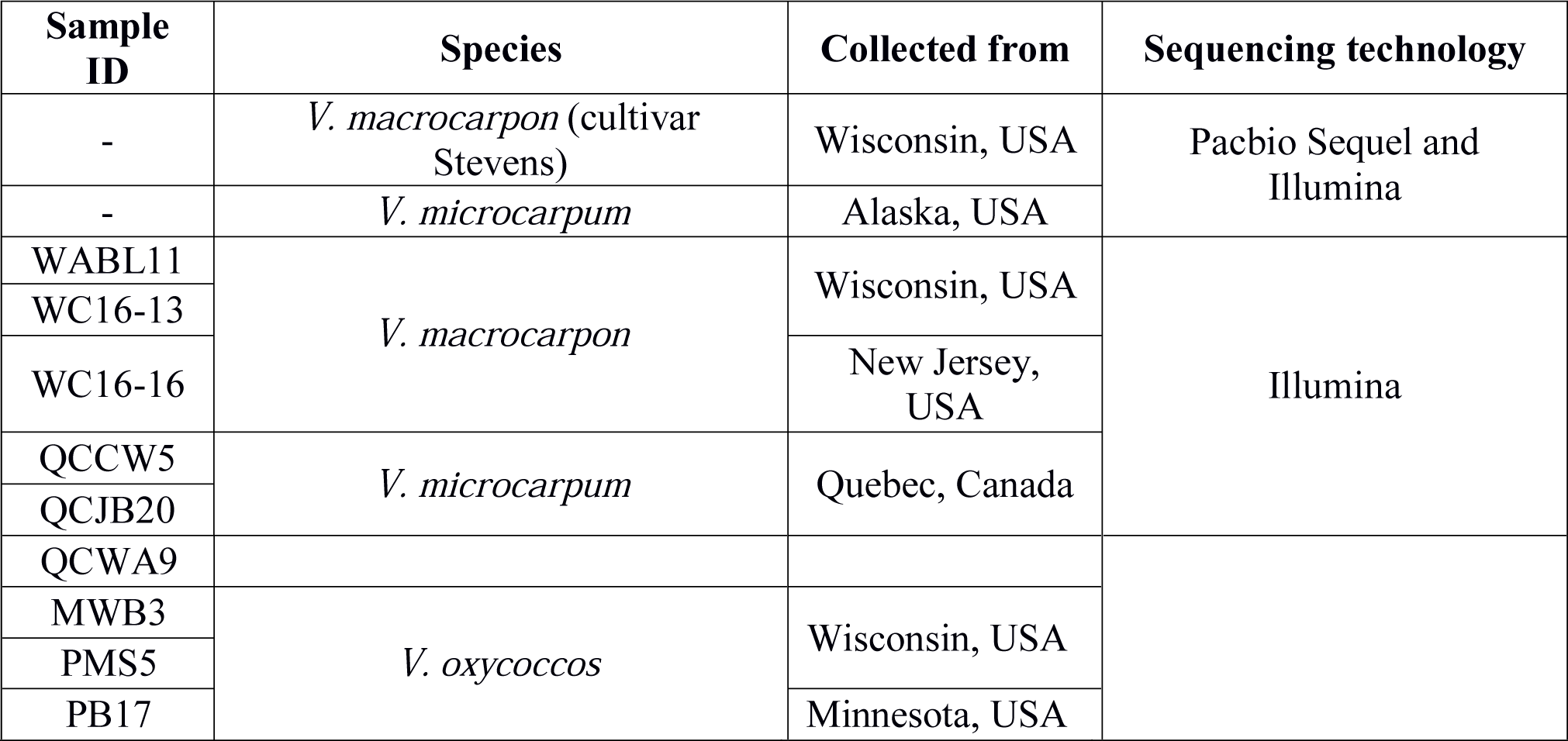
*Vaccinium* samples sequenced and assembled in this study

| Sample ID | Species | Collected from | Sequencing technology |
| --- | --- | --- | --- |
| - | *V. macrocarpon* (cultivar Stevens) | Wisconsin, USA | Pacbio Sequel and Illumina |
| - | *V. microcarpum* | Alaska, USA |  |
| WABL11 | *V. macrocarpon* | Wisconsin, USA | Illumina |
| WC16-13 |  |  |  |
| WC16-16 |  | New Jersey, USA |  |
| QCCW5 | *V. microcarpum* | Quebec, Canada |  |
| QCJB20 |  |  |  |
| QCWA9 |  |  |  |
| MWB3 | *V. oxycoccus* | Wisconsin, USA |  |
| PMS5 |  |  |  |
| PB17 |  | Minnesota, USA |  |

Genomic DNA was extracted using NucleoSpin® Plant II (Macherey-Nagel), and submitted to The Biotech Center, at the University of Wisconsin-Madison (Wisconsin, USA) for sequencing. Indexed samples were evenly loaded into 3 HiSeq 2500 Illumina lanes, 1×100bp. Illumina reads were cleaned using Trimmomatic [41] (minimum length 85bp and minimum quality 28), and then, mapped against our PacBio organelle genomes with bwa. Later, filtered reads were de novo assembled using abyss with the parameter k=64, and the contigs were ordered based the organelle reference genomes in Geneious R11. Variants were searched first, by aligning all 9 samples (global SNP across all 9 species), and then, by aligning samples of the same species (specific alignments, 3 sequences per species). Alignments were performed using the progressive method implemented in Mauve [42]; SNP were called after masking gaps in the alignment.

### 2.3. Phylogenetic analysis

Due to the recent increase in availability of plastid genomes, especially from the Ericales, we carried out a phylogenetic analysis using 68 genes (5 atp genes, ccsa, cema, matk, 11 ndh, 6 pet, 5 psa, 14 psb, rbcl, 7 rpl, 3 rpo, 11 rps, and 2 ycf), consistently present in 75 plastid genomes. In this analysis, 36 genomes corresponded to the Ericales. All genome sequences were downloaded from NCBI. Coding sequences were extracted in Geneious R11, and imported in R using the package Biostrings for filtering genes present in the majority of the species. Then, gene sequences were concatenated for each species, exported in fasta format, and imported back in Geneious R11 for sequence alignment and tree construction. Alignments were performed using the progressive method implemented in MAFFT v7 (Geneious R11 plugin) with the default parameters. Optimal trees were inferred using Maximum Likelihood as implemented in FastTree [44]. Since mitochondrial genomes of Ericales are still scarce, we did not performed any phylogenetic analysis based on mitochondrial sequences.

## 3. Results and Discussion

### 3.1. Cultivated and wild cranberries share identical organelle genomes

The first mitochondrial and plastid genomes in *V. macrocarpon* were published in 2014 [30] and 2013 [29], respectively. While these assemblies were highly reliable, the advantages of today’s single molecule real time sequencing allowed us to improve the current assemblies and generate novel organelle genomes of *V. microcarpum*.

The complete mitochondrial genome of *V. macrocarpon* obtained here was 468,115bp long, almost 10kb larger than the original version [30], while the plastid was 176,093bp long, just 48bp longer [29]. Both assemblies were resolved in a single circular scaffold with a mean coverage (based on PacBio corrected reads only) of 39.8X for the mitochondria, and 38.9X for the chloroplast. For the mitochondria, GC content was very similar with the previous version (45.3% versus 45.4%), whereas for the plastid, GC content was identical for both versions (36.8%). Moreover, GC content in both organelles fell within the expected range of the Asterids clade (based on the mitochondrial genomes for this group available in NCBI). To identify major structural changes between previous and current assemblies of V. macrocarpon organelles, we performed whole-genome alignments using mummer [45] (Figure 1). For the mitochondrial genome, 376 rearrangements were observed, from those, 50% corresponded to sequences smaller than 30 nucleotides and 81% to sequences smaller than 100 nucleotides. Although mummer identified rearrangements involving large sequences, they (and most of the smaller ones) shared perfect synteny between previous and current genome assemblies. For the plastid genome, 161 rearrangements were observed, from which 17% involved sequences smaller than 30 nucleotides and 43% smaller than 100 nucleotides. Similar to the mitochondrial genome, most of the rearrangements did not affect the overall synteny between both assemblies. Outputs from mummer are provided in the Supplementary File 1.

For *V. microcarpum* the genomes were resolved in a single circular scaffold with a mean coverage of 37.1% and 39.5% for mitochondrial and plastid, respectively. Remarkably, both mitochondrial and plastid genomes of *V. microcarpum* were completely identical to the ones in *V. macrocarpon*, with an exception of two synonymous single nucleotide polymorphisms (SNP) in the plastid, which are described below. Illumina resequencing of both *V. macrocarpon* and *V. microcarpum* confirmed our Pacbio assemblies.

Although uncommon, several studies in other plant species have reported the similarity of organelle genomes among cultivated and wild species. For example, the mitochondrial genomes of wild (*Hordeum vulgare* ssp. spontaneum) and cultivated (*H. vulgare* ssp. vulgare) barley, which has a history of 10,000 years after domesticated from the wild progenitor, are identical with the exception of three SNP [46]. Similarly, a study regarding plastid inheritance in *Nicotiana tabacum* found very few SNP when comparing the later with *N. sylvestris*, its maternal wild progenitor [47].

**Figure 2.**
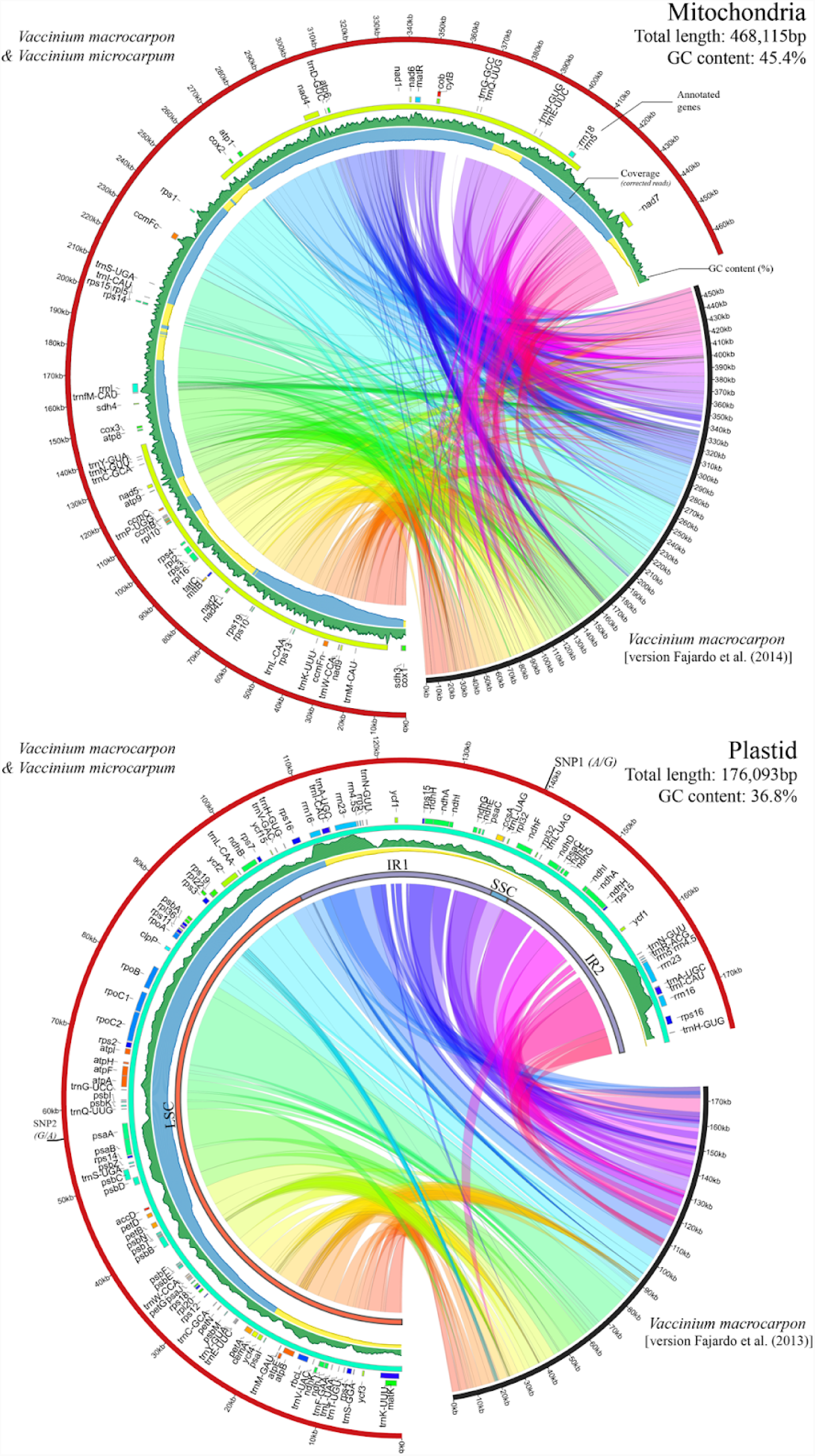
Organelle genomes of *V. macrocarpon* and *V. microcarpum*. A comparison between the PacBio assemblies presented in this study and previously published cranberry organelle versions [29,30] are shown; collinear regions were calculated using mummer with default parameters (see Materials and Methods). Since *V. macrocarpon* and *V. microcarpum* presented near identical organelles, for the plastid genome, only SNP positions are provided and the mitochondria is shown as identical. Gene annotations are shown as tiles in internal tracks; colors were randomly assigned to highlight different categories of protein-coding genes, ribosomal RNA, and tRNA. Sequencing coverage is provided in area plots oriented inwards, where blue, yellow, and red, represent genome areas with >30X, >10X, and ≤10X coverage, respectively (based on PacBio corrected reads). GC content is also provided as a green-colored area plot. In the plastid, large single copy (LSC), small single copy (SSC), and inverted repeats (IR1 and IR2) are highlighted with red, blue, and purple concentric bars.

### 3.2. Genome architecture and gene content

Since *V. macrocarpon* and *V. microcarpum* contained near identical organelle genomes, the following results regarding gene content, repeated sequences and other statistics, apply for both species.

For the mitochondria, gene content was similar to the previous genome version [30]. We found 9 genes of the respiratory chain complex I (nad-NADH dehydrogenase), 2 of the complex II [sdh-succinate oxido-reductase, one more than in Fajardo et al. (2014)], one of the complex III (apocytochrome b), 3 of the complex IV (cox-cytochrome oxidase), and 5 of the complex V [atp-ATP synthase, one more than in [30]]. In addition, we found 4 genes of the cytochrome c biogenesis (ccm), two for the cytochrome b (cytB and petL), one transcription gene (mat-maturase), one transporter protein (mtt-B), 3 ribosomal RNA (rrm), 13 ribosomal proteins (rpl and rps genes) and a malonate uptake protein (tatC). In total, we found 17 tRNA, from which 3 were duplicated (trnL, trnM, and trnS). In Fajardo et al. (2014), the authors reported the presence of two copies of tRNA-Sec, an unusual transfer RNA in land plants; here, we confirm its presence, but in a single copy. [exon 1 of copy 1 (located at ∼45kb) has a single different nucleotide]. Out of the 42 protein-coding genes found in the mitochondrial genome, 3 contained non ATG start codon, and 8 had undetermined stop codons (different than TAA, TGA, or TAG). Among these, the gene petL had both start and stop codons undetermined, as well as it was 73 nucleotides long. Similarly, gene mttB was expected to have non ATG start codon based on previous studies [48–51]. Protein-coding genes varied largely in size and intervals, ranging from a few hundreds of nucleotides and a single interval (one exon), to thousands of nucleotides and up to 5 intervals (i.e. *nad7* and *nad2*). Regarding SSR, we observed 1449 motifs with total lengths ranging from 6 to 26 nucleotides. As expected, dinucleotides and trinucleotides were the most abundant (80.0% and 15.6%, respectively); tetra, penta and hexanucleotides account for less than 5%. The repeat AG was the most abundant motif among the dinucleotides. A complete list of repeated sequences in the mitochondria is provided in Supplementary File 1.

Most of the genes found in the plastid were also consistent with the previous version published by [29]. The annotated sequences were classified as follows: out of the 71 unique protein-coding genes, 6 corresponded to ATP synthase subunits (atp genes), 10 NADH-dehydrogenase subunits (ndh genes), 6 cytochrome b/f complex subunits (pet genes), 5 photosystem I subunits (psa genes), 16 photosystem II subunits (psb genes), 8 large ribosomal protein units (rpl genes), 12 small ribosomal protein units (rps genes), and 4 RNA polymerase subunits (rpo genes). Other important genes such accD, ccsA, infA, matK, rbcL were also found. More than 95% of these genes showed regular start and stop codons. Five hypothetical genes (*ycf*) were found, although only one showed standard start and stop codons. In addition, 4 ribosomal RNA and 27 unique transfer RNA were found. The length of the large single copy (LSC), small single copy (SSC), and inverted repeats (IR) fragments were 104,592bp, 3,025bp, and 34,238bp, respectively. All ribosomal RNA were located in the IR, therefore two copies of each gene were found; on the other hand, *ndhF* was the only gene in the SSC. Only 492 SSR were found in the plastid genome with the following proportions: dinucleotide 77.2%, trinucleotide 13.4%, tetranucleotide 5.9%, and both penta and hexanucleotide with less than 2.0% each. The dinucleotide motif AT was the most abundant (Supplementary File 1). SSR frequency and distribution of SSR motif type were different compared with previous studies in cranberry [33]. In the previous study, the mitochondria had fewer SSR (1 every 3.8kb) than the plastid (1 every 2.0kb) genome (in average), and the tetranucleotide motifs were the predominant for both organelle genomes.

As mentioned before, both mitochondrial and plastid genomes were virtually identical between *V. macrocarpon* and *V. microcarpum*. Only two SNP were found in the plastid; the first one (G in *V. microcarpum*, A in *V. macrocarpon*, at position 139,582bp), located in the ndhF (at the SSC fragment), was a non-synonymous mutation producing isoleucine in *V. macrocarpon* and valine in *V. microcarpum*, two similar non-polar aminoacids. The second SNP (A in *V. microcarpum*, G in *V. macrocarpon*, at position 57,591bp) was located in the psaA gene, and it resulted in a synonymous change (translate to the aminoacid glycine in both cases). Pacbio genome assemblies can be provided under request.

### 3.3. Phylogenetic analysis of *V. macrocarpon* and related species

To study the phylogenetic position of *V. macrocarpon* within the Ericales, we performed an analysis based on 68 genes consistently present among 36 Ericales genomes, and 39 from other families (Figure 3). As expected, the Ericales formed a monophyletic group. The topology of our tree mirrors relationships recovered from a more comprehensive analysis of a supermatrix that includes nearly 5000 species and 25 loci [52]. In both analyses, the balsaminoids (represented by *Impatiens piufanensis* and *Hydrocera triflora* in our data) are placed sister to the rest of the Ericales; and the Ericaceae are sister to the sarracenioids (represented by *Actinidia* in our data) in a strongly supported clade. Our data included two additional Ericaceae, *Arbutus unedo* and *Chamaedaphne calyculata.* Of these, *V. macrocarpon* was most closely related to *C. calyculata. Chamaedaphne calyculata,* a circumboreal species with a recently published plastid genome [53], shares almost an identical genome architecture with *V. macrocarpon*, with the most profound changes in the inverted repeat regions. Moreover, *C. calyculata* shares many phenotypic traits with wild *V. macrocarpon* plants [54].

**Figure 3.**
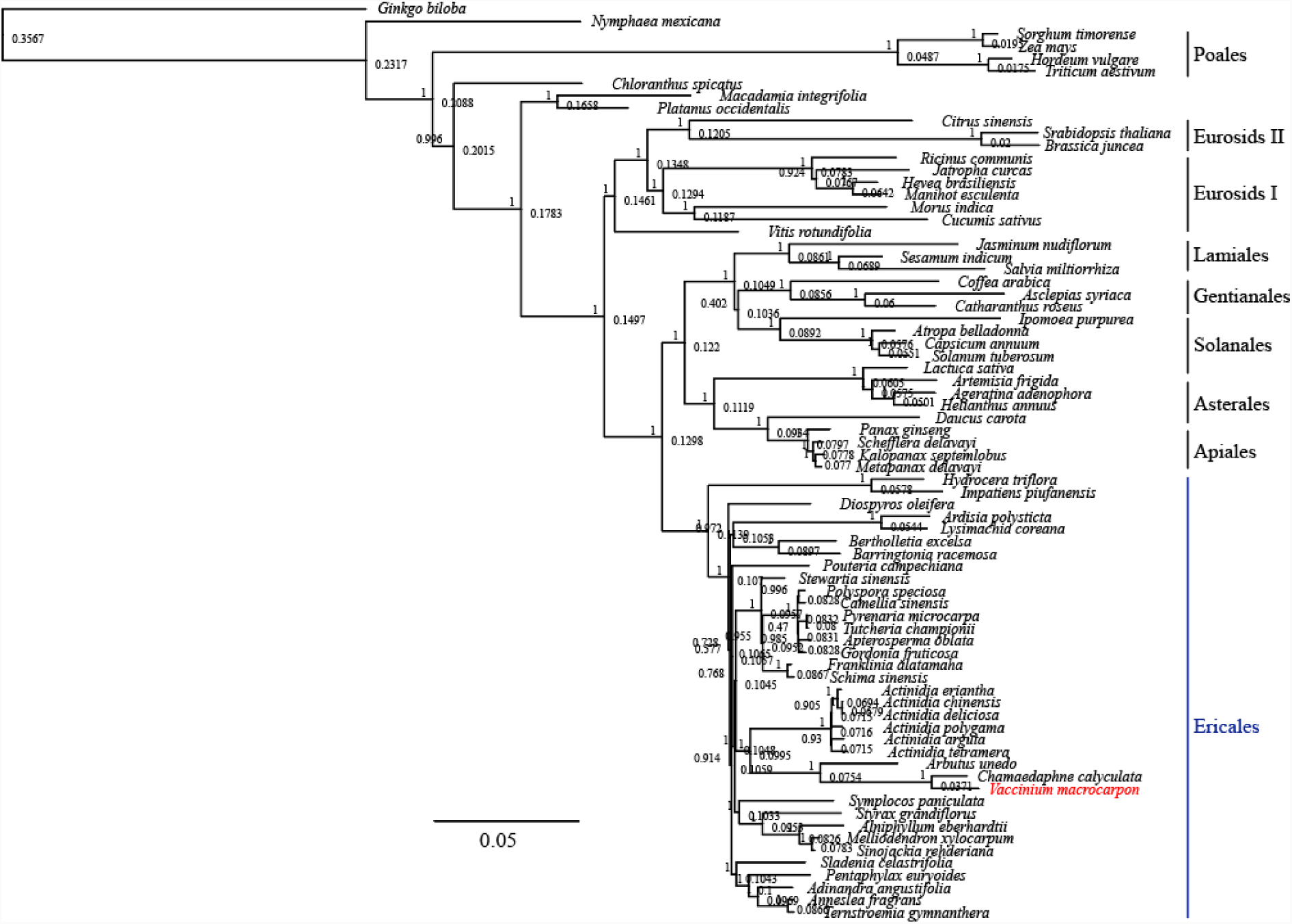
Phylogenetic relationship of *V. macrocarpon* (and Alaskan *V. microcarpum*) with related species based on 68 protein-coding genes shared by all plastid genomes. Tree constructed in FastTree using the Generalized Time-Reversible (GTR) model. Branch labels correspond to the FastTree support values, whereas node labels show node heights. *Ginkgo biloba* was used as outgroup.

### 3.4. Divergent *V. microcarpum* can be found in eastern Canada

Due to the remarkable similarity found between *V. macrocarpon* and *V. microcarpum* organelle genomes, we expanded our study by resequencing additional accessions from a broader geographic area. In particular, we included samples of *V. microcarpum* collected in Quebec Canada, *V. oxycoccos* (4x) collected in Wisconsin and Minnesota, USA, as well as *V. macrocarpon* from Wisconsin and the east coast of USA (Table 1). For these samples, Illumina resequencing allowed us to partially assemble both mitochondrial and plastid genomes at a very high coverage (>1000X). After removing contigs smaller than 5kb, mitochondrial genome assemblies for all 9 samples had a length between 394kb and 428kb distributed among 27 to 30 scaffolds (additional statistics regarding assemblies are provided in Supplementary File 1). For the plastid, assembly lengths (taking in account only one of the inverted repeats) ranged from 76kb to 95kb among 7 to 10 contigs. The incompleteness of these de novo assemblies could have been produced by two main causes. First, the use of very short reads (1×100bp) could have hampered the assembly process. And second, since these *de novo* assemblies were conducted using a subset of raw reads that mapped to our *V. macrocarpon* PacBio assembly, potential novel sequences absent in either the reference or the additional accession could have been missed.

After orienting and ordering the plastid and mitochondrial contigs for all 9 samples (based on *V. macrocarpon*), we aligned them and identified SNP variants. By considering aligned and ungapped regions involving all 9 samples (using our *V. macrocarpon* PacBio assemblies as reference), we discovered 831 SNP in the mitochondria (Figure 4A and Table 2) and 121 in the plastid (Figure 4B and Table 2). Of the SNP found in the mitochondrial genome, 4 were located in the *nad1* gene, 15 in the *nad2* gene, and 1 in the *rps14* gene. In the plastid, 39 SNP were found in 13 different genes, being *ycf2* the one with the most (11). A large portion of SNP were found in the regions delimiting the inverted repeat regions, which was expected given their volatility [55]. When aligning only the wild *V. macrocarpon* samples, 10 SNP were found in the mitochondria and 11 in the plastid (Figure 4C, top panel). Similarly, alignments of only the tetraploid *V. oxycoccos* resulted in 11 SNP each for both mitochondria and plastid (Figure 4C, bottom panel). In Contrast, the alignment of the Canadian *V. microcarpum* samples resulted in 232 SNP in the mitochondria and 332 SNP in the plastid; from these, more than 95% of the SNP were called because sample QCWA9 showed alternative alleles (Figure 4C, middle panel, for space reasons, only first alignment position are shown).

**Figure 4.**
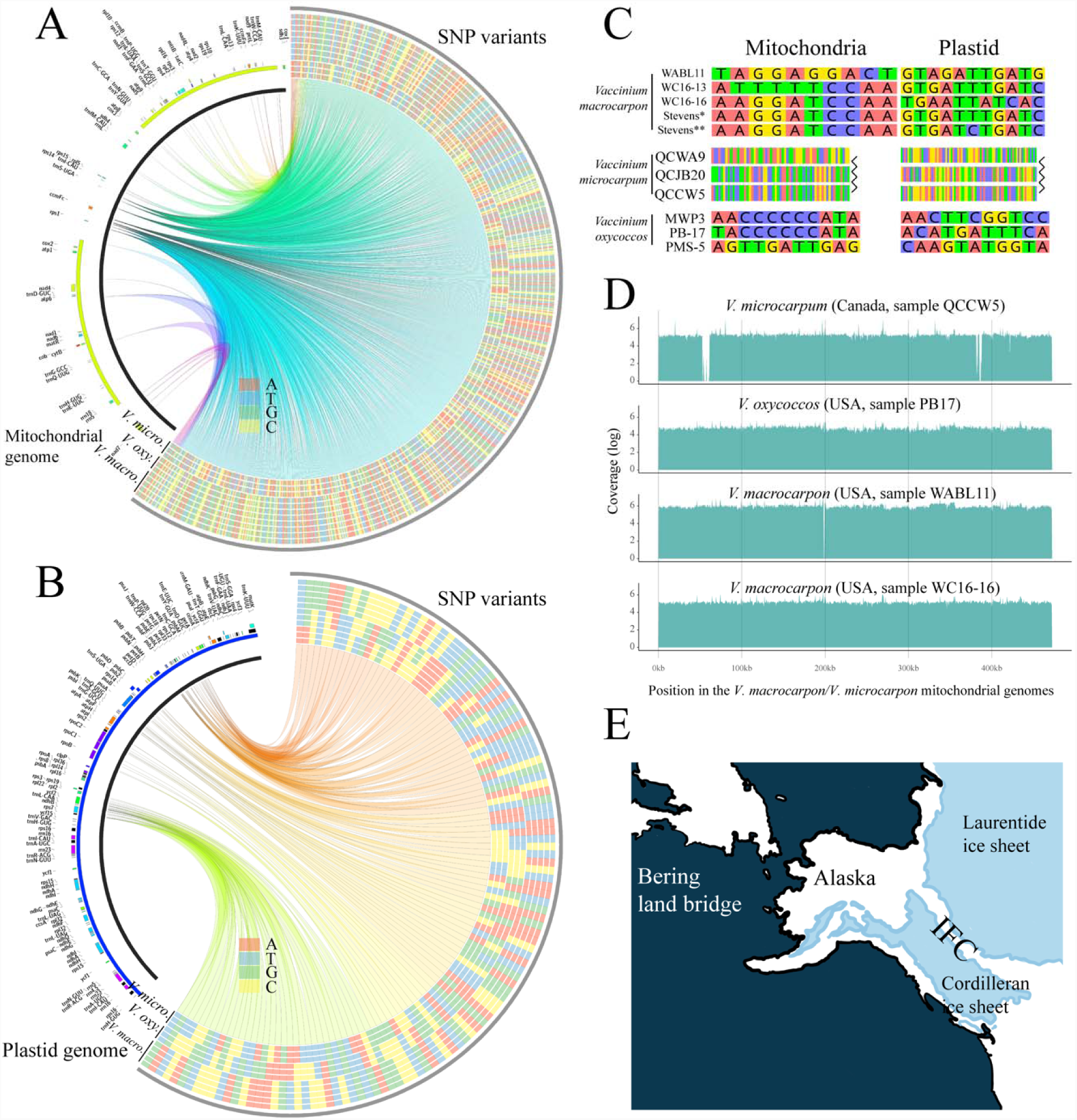
SNP variants among *V. macrocarpon, V. microcarpum* and *V. oxycoccos*. SNP variants detected in mitochondria (AB) and plastid (BC) genomes among all 11 samples sequenced with both Illumina and Pacbio. On the heatmaps (right arcs), the inner three rings correspond to the *V. microcarpum* samples QCCW5, QCJB20, and QCWA9; the next three rings (4-6) correspond to the *V. oxycoccos* samples MWPB3, PB17, and PMS5; rings 7-9 correspond to the wild *V. macrocarpon* samples WABL11, WC16-13, and WC16-16; and rings 10-11 correspond to cultivated *V. macrocarpon* sequenced with Illumina and Pacbio, respectively (note these last two sequences were identical to the Alaskan *V. microcarpum* sequences). Each SNP is connected by a colored link to its physical position within either the mitochondria or plastid genome (leftright arcs); link colors are meaningless. For reference purposes, gene locations were added on the left arcs and colored by category. (CD) SNP variants called when aligning sequences by species; for the *V. macrocarpon* alignment, Stevens* correspond to the assembly carried out with Illumina data (used to validate the PacBio assembly), whereas Stevens** correspond to the one with PacBio; for *V. microcarpum*, only the first positions of the alignment are shown. (D) Coverage plots using Illumina filtered raw reads and *V. macrocarpon*/*V. microcarpum* mitochondrial genomes as reference (additional coverage plots are provided in Figures S1 and S2). (E) Ice extent between 9-10k years ago showing the ice free corridor (IFC).

**Table 2.**
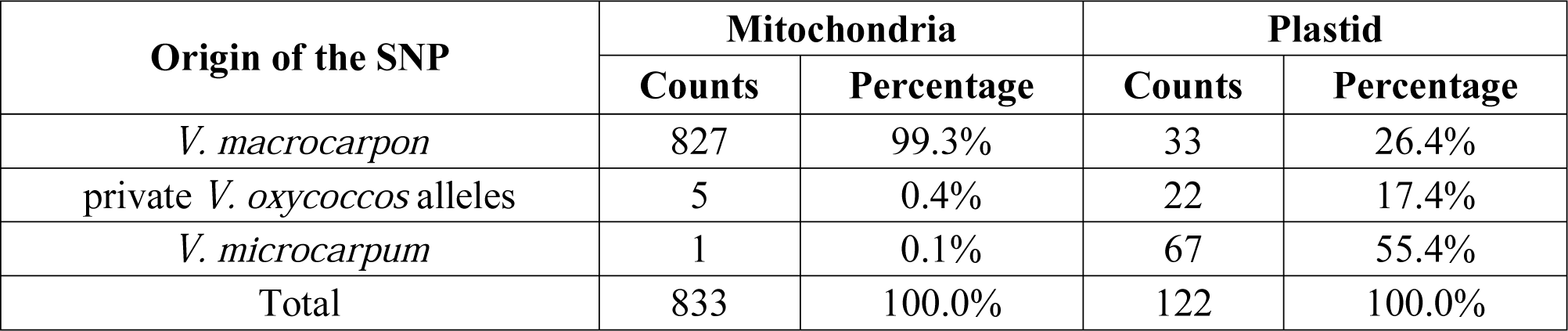
Proportion of *V. oxycoccos* SNP found in either *V. macrocarpon, V. microcarpum*, or that represent novel alleles.

Additionally, we carried out an alternative approach in which, instead comparing the de novo assemblies, the Illumina raw reads (previously trimmed and cleaned) were directly mapped using our *V. macrocarpon* as reference and low coverage regions were recorded. This strategy revealed that mitochondrial genomes of the wild *V. macrocarpon* samples WC16-13 and WC16-16 were very similar to our reference; however, we could not find Illumina reads of WABL11 to map a ∼1kb region at position ∼198kb of our *V. macrocarpon* assembly. Also, when mapping mitochondrial raw reads of our Canadian *V. microcarpum* samples, we observed a region of ∼8kb at position ∼55kb, and a two-interval region of ∼7kb at position ∼384kb that we could not find Illumina reads to map. Interestingly, all the *V. oxycoccos* (4x) samples showed a continuous and ungapped coverage along both plastid and mitochondrial genomes. Figure 4D shows the unique coverage plots observed across all nine samples in the mitochondria; in Figures S1 and S2, all coverage maps for both mitochondria and plastid genomes are provided.

### 3.5. Relationships among diploids, and the link between Alaskan *V. microcarpum* and *V. macrocarpon*

*Vaccinium macrocarpon* and *V. microcarpum* displayed dramatically different genetic diversity. Our samples of *V. macrocarpon* span 1700 km from Wisconsin to the east coast. Despite the geographic distance, four samples yielded only 21 intraspecific SNPs combined for the chloroplast and mitochondrial genomes. In contrast, we found 564 intraspecifc SNPs in three *V. microcarpum* samples collected within 260 km along the James Bay Coast. This is a very modest representation of a species with a circumboreal distribution; more comprehensive sampling is likely to reveal further diversity.

The organelle genomes shared by the Alaskan *V. microcarpum* sample and all the *V. macrocarpon* samples is suprising. Given the unexpected diversity in our *V. microcarpum* plants, this could be explained by the persistence of multiple ancestral cytoplasmic lineages across the range of *V. microcarpum*, including the one now present in *V. macrocarpon*. In other words, the shared genomes are the consequence of incomplete lineage sorting among *V. macrocarpon* and *V. microcarpum.*

Alternatively, the shared genomes could be the result of past gene-flow between the Alaskan *V. microcarpum* populations and *V. macrocarpon* in eastern North America. Our *V. macrocarpon* accession was collected in southern Alaska, in the region of the ice-free corridor (IFC) that connected eastern Alaska with the north-central region of the US during the deglaciation period between 8,000 and 11,000 years (Figure 4E) [56]. The IFC enabled the movement of humans and wildlife across the continent thousands of years before the Laurentide ice sheet retreated from eastern Canada, which could explain a closer genetic relationship between *V. macrocarpon* and the Alaskan *V. microcarpum* than the geographically closer populations in Quebec. Further sampling across the range of *V. microcarpum* is needed to clarify this situation.

### 3.6. The origin of the tetraploid *V. oxycoccos*

Different studies have discussed the tetraploid origin of *V. oxycoccos*, but no definitive answer has been provided. Two main theories have been proposed. The first one describes that *V. oxycoccos* arose as a hybrid of the two diploid taxa, *V. macrocarpon* and *V. microcarpum* ([18], see above), whereas the second suggest that *V. oxycoccos* is most likely the autopolyploid descendant of the diploid *V. microcarpum* [19]. Since *V. oxycoccos* was included in our resequencing analysis, the SNPs found across all three species may provide evidence about the origin of *V. oxycoccos*.

For 825 of the 831 mitochondrial SNPs (99.3%), *V. macrocarpon* and *V. oxycoccos* shared the same allele, with the alternative allele present in *V. microcarpum* (Table 2). *Vaccinium oxycoccos* and *V. microcarpum* shared the same allele at a single mitochondrial SNP (< 1%). This strong link to *V. macrocarpon* refutes the hypothesis that *V. oxycoccos* is an autopolyploid descendant of *V. microcarpum*. The simplest explanation is that *V. oxycoccos* is an allotetraploid, and that *V. macrocarpon* was the ancestral cytoplasmic donor.

However, the chloroplast data challenges this simple explanation, as the *V. oxycoccos* genome appears to contain a mixture alleles from both diploid species. *Vaccinium macrocarpon* and *V. microcarpum* are distinguished by 99 chloroplast SNPs. Of these, *V. oxycoccos* had the “*microcarpum*” allele for 67 SNPs, and the “*macrocarpon*” allele at 32 SNPs. We are aware of no mechanism that would enable recombination between chloroplast genomes from diploid ancestors to yield a mixture of genomes in the derived polyploid. This may indicate that our sampling was not adequate to capture the full scope of variation in the chloroplast genomes among the three species. That is, the alleles that are unique to *V. macrocarpon* and *V. microcarpum* in our data may actually be present in both species. Alternatively, a third diploid species may have been involved in the formation of tetraploid *V. oxycoccos.*

The possible existence of undocumented genetic diversity is further supported by the contrasting results from the PacBio sequencing. These data show that the Alaskan *V. microcarpum* sample possesses mitochondrial and chloroplast genomes identical to *V. macrocarpon*, and markedly different from the genomes of conspecific samples from Quebec. This could be explained by the preservation of multiple ancestral organelle lineages in the circumboreal *V. microcarpum*, including the one present in more geographically restricted *V. macrocarpon.*

Alternatively, it could be an indication of cryptic variation in the *V. oxycoccos/V. microcarpum* group. Porsild (1938) recognized a western cranberry species (*Oxycoccos ovalifolius* (Michx.) Porsild). Later authors have subsumed it within *V. oxycoccos* (along with *V. microcarpum*, [5]). However, genetic studies of *V. microcarpum* have been geographically limited: Mahy et al. (2000) included only Alaskan *V. microcarpum* samples, and Smith et al. (2015) was restricted to western Quebec. The results of the current study suggest that a broader assessment of genetic diversity in diploid cranberries across their northern range may reveal taxonomically and agronomically valuable variation.

### 3.7. Conclusions and further directions

Our study of the organelle genome architecture of *V. macrocarpon, V. microcarpum*, and *V. oxycoccos*, and their diversity across North America, revealed a remarkable similarity between the cytoplasms of Alaskan *V. microcarpum* and cultivated *V. macrocarpon*. Moreover, we showed that a highly differentiated *V. microcarpum* can be found in eastern Canada, and suggested that the spatial and temporal dynamics of the last glaciation period could had shaped the genetic diversity among cranberry species. Further studies should investigate the genomic rearrangements observed in Canada’s *V. microcarpum* samples as well as elaborate a more extensive sampling and genomic profiling of germplasm across Alaska and the “ice-free corridor”. In summary, this study provided important insights about the movement and genetic differentiation of cranberry and its wild relatives in North America, which is of vital importance for the inclusion of wild germplasm into cranberry breeding programs.

## Author contributions

LDG, LRB and JZ design the study; TS and JZ provided plant material; LDG and LRB performed the experiment; LDG analyzed the data; LDG, LRB, TS and JZ wrote the manuscript.

## Funding statement

This project was supported by USDA-ARS (project no. 5090-21220-004-00-D provided to JZ); WI-DATCP (SCBG Project #14-002); Ocean Spray Cranberries, Inc.; Wisconsin Cranberry Growers Association; Cranberry Institute. LDG was supported by the Consejo Nacional de Ciencia y Tecnología (Mexico) and the Gabelman-Seminis Graduate Fellowship. LRB was supported by the UW Madison SciMed GRS.

## Acknowledgments

We are grateful to Drs. Mike Havey and Sean Schoville at UW-Madison for their helpful insight. Also we want to thank the Wisconsin and Minnesota Department of Natural resources for the permits to collect in their lands as well as the US Forest Service. JZ would like to express his gratitude through Ps 136:1. The authors also thank the anonymous reviewers who helped enhance the quality of this paper.

