## Supplementary figures and images for "Pacbio sequencing reveals identical organelle genomes between American cranberry (*Vaccinium macrocarpon* Ait.) and a wild relative"

### Figure S1

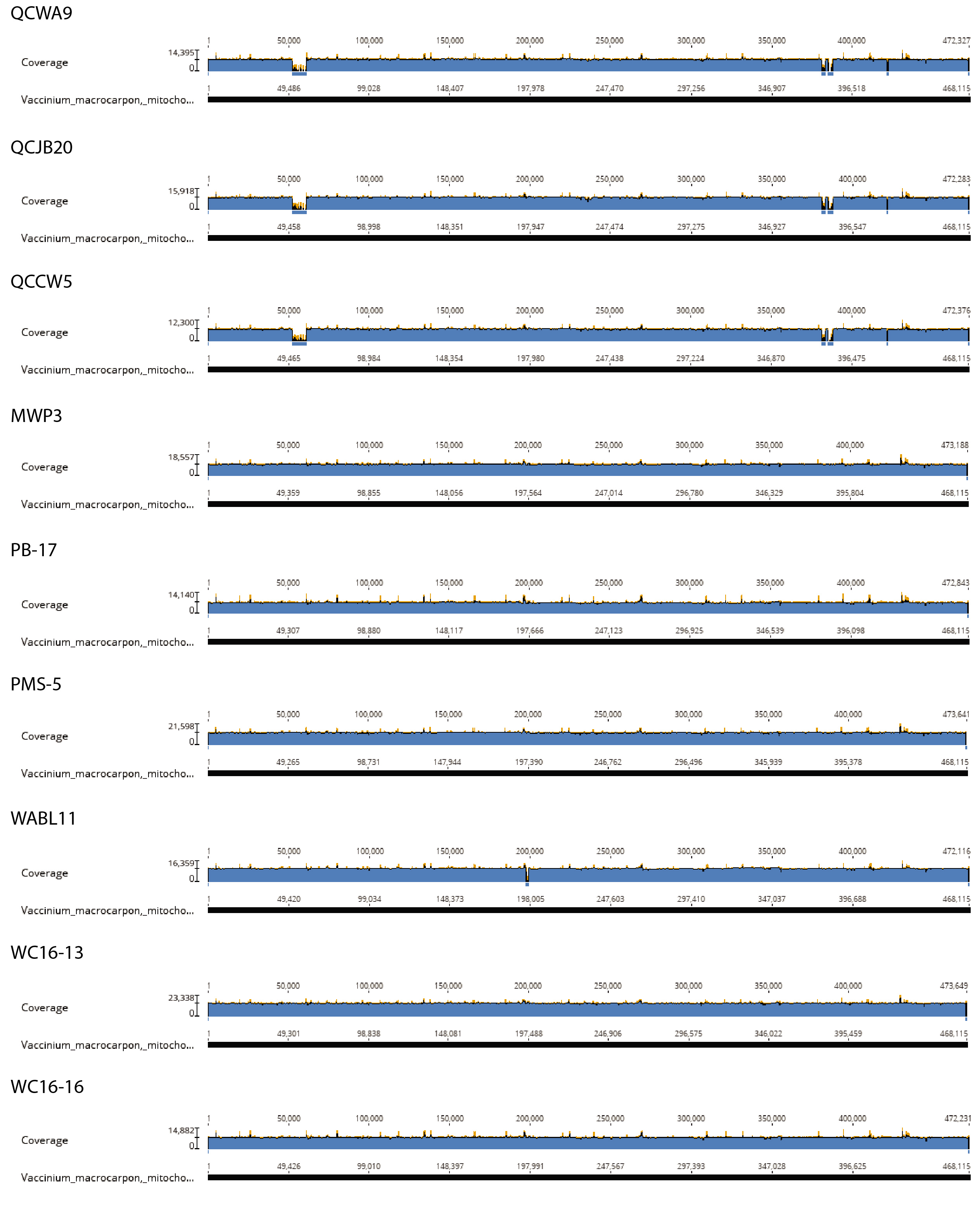

### Figure S2

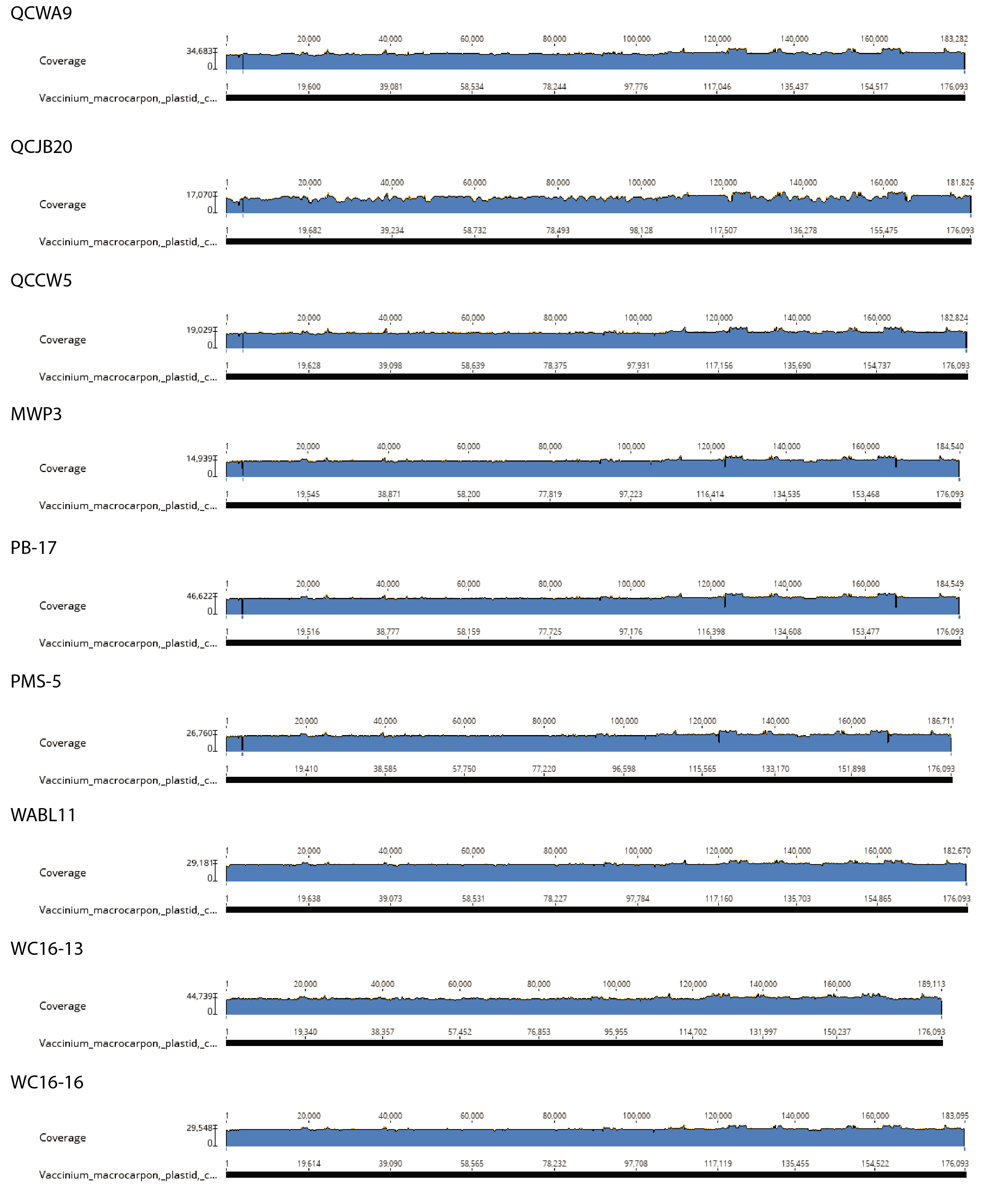
